# Deep Sound Synthesis Reveals Novel Category-Defining Sound Features in the Human Auditory Cortex

**DOI:** 10.1101/2025.02.14.638230

**Authors:** Lidongsheng Xing, Elia Formisano, Lars Riecke

## Abstract

The human auditory system extracts meaning from the environment by transforming acoustic input signals into semantic categories. Specific acoustic features give rise to distinct categorical percepts, such as speech or music, and to spatially distinct preferential responses in the auditory cortex. These responses contain category-relevant information, yet their representational level and role within the acoustic-to-semantic transformation process remain unclear. We combined neuroimaging, a deep neural network, a brain-based sound synthesis, and psychophysics to identify the sound features that are internally represented in the speech- and music-selective human auditory cortex and test their functional role in sound categorization. We found that the synthetized sounds exhibit unnatural features distinct from those normally associated with speech and music, yet they elicit categorical cortical and behavioral responses resembling those of natural speech and music. Our findings provide new insights into the fundamental sound features underlying speech and music categorization in the human auditory cortex.

## Introduction

Speech and music are two of the most fundamental communicative signals for humans. They emerge from different natural sound sources (e.g., human voice or musical instrument) [1,2] and consequently differ in their characteristic acoustic properties (e.g., temporal or spectrotemporal modulations [3,4]). Speech [5–7] and music [6,7] stimuli elicit preferential responses in distinct, partially overlapping regions of the non-primary auditory cortex [7–9] with some lateralization [2,10,11] (speech: left hemisphere, music: right hemisphere), suggesting they are processed by specialized mechanisms. Previous studies [6,11,12] have attributed these category-preferring responses to the distinct acoustic properties and/or perceptual attributes of speech and music. For example, Albouy et al (2020) demonstrated the relevance of spectrotemporal modulation rates, whereas Norman-Haigner et al (2015) emphasized the role of the speech- and music-category percepts beyond acoustic features such as frequency or spectrotemporal modulation. Based on these findings, speech- and music-preferential cortical processing might emerge from the acoustic features and/or the categorical percepts that speech and music induce.

Given the still limited number of studies addressing this topic, a large number of potentially relevant sound features for cortical speech/music processing remains unexplored. In particular, it is unclear how category-preferring cortical regions internally represent the preferred sound. By “internal sound representation” we refer to (the level of abstraction of) category-related information that is primarily processed within the region, which could range from basic fine-grained spectrotemporal information to abstract semantic labels.

A conventional way of addressing these questions involves an exhaustive testing of a large set of pre-defined, experimentally controlled sound features hypothesized to contribute to categorization. A more efficient and powerful way may be to identify possibly novel, categorization-relevant sound features within the vast feature space of an artificial deep neural network (ANN) trained to perform sound classification. The latter ANN transforms an acoustic input signal into a sound-category label across several hierarchical processing stages (i.e., layers), where consecutive layers represent the original acoustic input as a combination of increasingly abstract features. The representation of natural sounds in such an ANN has been shown to correlate with the responses that these sounds evoke in the human auditory cortex [13,14], implying that the ANN may be utilized to predict auditory cortical response to natural sounds. Indeed, its ability to represent sounds at various abstraction levels and to predict cortical responses to these sounds make the ANN potentially suitable for identifying the internal sound representation of a given cortical region from the ANN layers. In this feature-identification approach, the ANN’s predictive power is exploited to determine stimuli that are most effective in eliciting a response in a given brain region (according to the ANN’s prediction), and the features of those stimuli can be inferred to contain the information that is primarily processed within that region. Application of this approach in monkey electrophysiology and human neuroimaging studies has yielded novel insights into stimulus features underlying specific brain functions. For instance, human functional magnetic resonance imaging (fMRI) studies have determined visual objects [15] and sentences [16] that an ANN predicted to maximally activate category-selective regions in the visual cortex and cortical language network, respectively. Moreover, monkey electrophysiology found novel, ANN-generated visual stimulus features that activate visual cortical neurons [17] even to higher levels than natural stimulus features do [18], and demonstrated the tuning of so-called cortical “face cells” to non-facial stimulus features [19].

In the present study, we applied the ANN-based feature-identification approach to inform fMRI and behavioral experiments aimed at identifying the fundamental sound features underlying sound categorization in the human auditory system. We first established a linear mapping from representations of natural categorical sounds (as modeled by individual ANN layers) onto fMRI responses to the same sounds (recorded from speech- and music-preferring auditory cortical regions, ROIs). Utilizing this mapping in a gradient-based sound synthesis, we obtained novel auditory stimuli predicted to strongly activate the speech- or music-ROI. In subsequent in-silico, fMRI, and behavioral experiments, we characterized the categorical features of these novel stimuli and tested their relevance for speech- and music categorization in the human auditory system.

## Results

Fig 1 renders a schematic overview of the approach that we applied individually to each participant.

**Fig 1.**
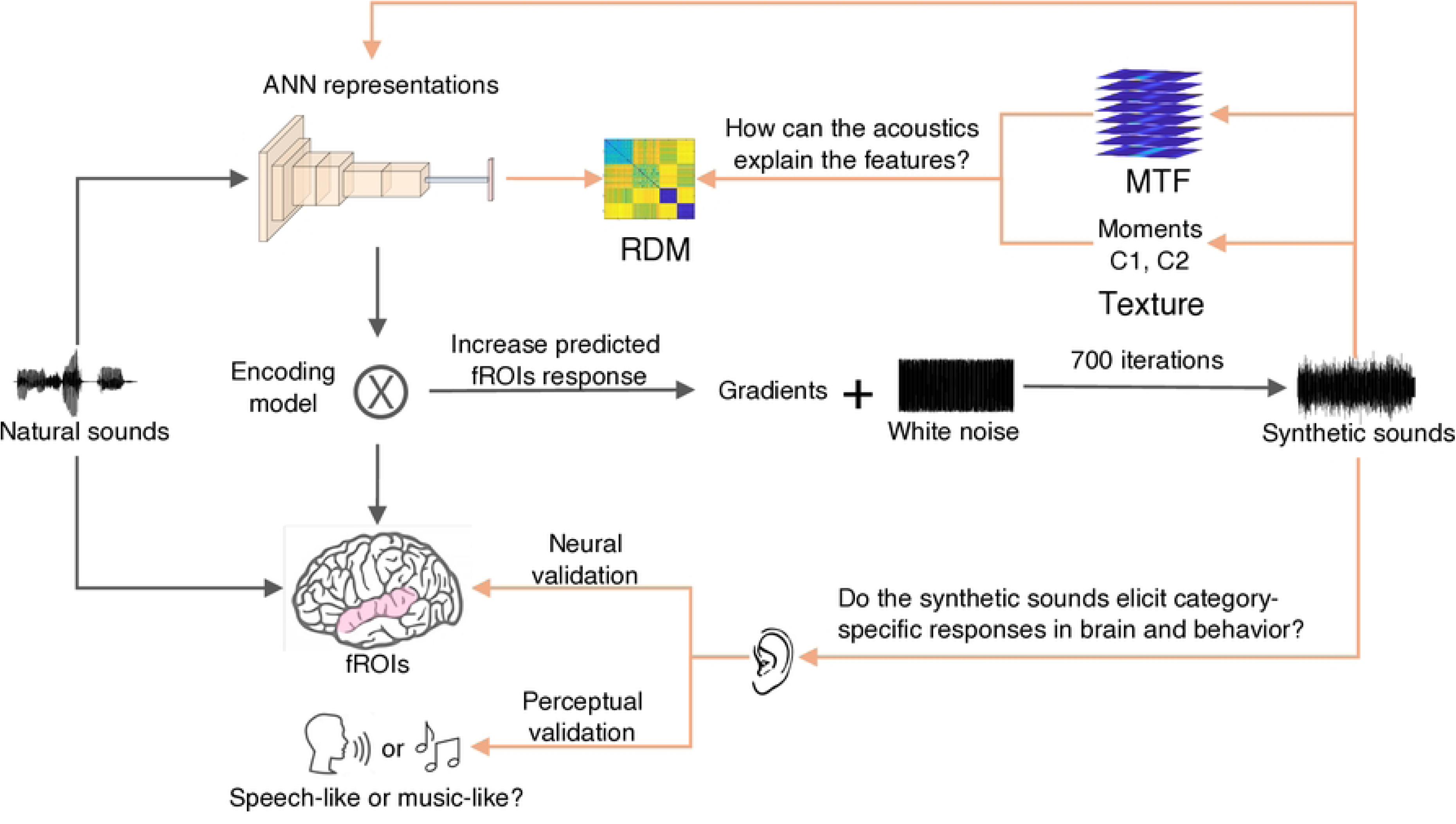
Overview of the approach. Black arrows indicate steps involved in the ANN-based sound feature identification. Orange arrows indicate steps involved in the characterization of the identified features and the testing of their putative functional role in sound categorization. In the first part, natural categorical sounds were presented to both ANN and participants, and the ANN and fMRI responses to these sounds were recorded. A linear encoding model was fitted that mapped the ANN responses onto the fRMI responses within speech and music fROIs in the auditory cortex. Utilizing this mapping and a gradient-based approach, features embedded in the ANN that maximized the predicted response of the speech fROI or music fROI were sonified, resulting in novel synthetic sounds. In the subsequent experimental part, the individually synthetized sounds were presented to both ANN and the respective participants, and the ANN responses, fMRI responses, and behavioral responses in a sound-categorization task were recorded. To investigate the categorical nature and uniqueness of these features, ANN responses to the synthetic sounds were compared to ANN responses to natural speech or music (using an analysis of representational dissimilarity matrices, RDM) and their similarity to acoustic features known to distinguish sound categories (MTF and acoustic texture) was assessed. To test the functional relevance of the novel sound features for sound categorization, participants’ fROI and behavioral responses to sounds synthesized to activate the speech fROI were compared vs those synthesized to activate the music fROI.

### Prediction of auditory cortical responses from ANN responses to natural sounds

We presented a set of natural sounds to ANN and participants, and recorded the responses of the convolutional layers and participants’ cortex. The ANN used here was the VGGish, which possesses a relatively simple architecture [20] and has been shown to accurately predict human auditory cortical fRMI responses to natural sounds [14]. We identified speech- and music-preferring voxels in the auditory cortex as follows: first, we selected voxels showing significant responses to sounds compared to a baseline without any natural sound stimulation (p < 0.001, uncorrected). Second, from this subset of sound-responsive voxels, we further selected voxels whose response profile to all natural sounds showed high reliability across fMRI sessions (see Methods). Finally, to identify the speech- and music-fROIs, we submitted the retained voxels to a statistical analysis contrasting speech vs. non-speech and music vs. non-music, and selected the top 5% of voxels (based on highest t-value, all p < 10^-4^; number of voxels min: 26, max: 110, median: 50.5 across participants). For the ANN responses, we selected four evenly distributed layers from the VGGish architecture (conv1, conv3_1, conv4_1, fc1_1; referred to as “c1, c3, c4, f1”), which we mapped onto the overall, spatially averaged fROI responses (see Methods). Fig 2 presents the results of the participant who showed the strongest effects (P2; for other participants, see S1 Fig). Fig 2A shows the cortical location of the speech- and music-fROIs, which occupied multiple portions of the STG in both hemispheres. Fig 2B shows the overall response of the fROIs to natural sounds. Per fROI definition, natural speech- and music-evoked responses were larger than zero (baseline) and the overall responses to the respective other natural sound categories. Fig 2C shows the prediction accuracy of the encoding model. Each layer could predict each fROI’s overall response to the natural sounds with an accuracy significantly above chance level (estimated from 1000 permutations, p = 0.001, corrected across layers). For each fROI, the best and poorest prediction was achieved respectively with layer 4 (median proportion explained variance in speech fROI: *R^2^* = 0.70, music fROI: *R^2^* = 0.50) and layer 1 (speech fROI: *R^2^*= 0.54, music fROI: *R^2^*= 0.36).

**Fig 2.**
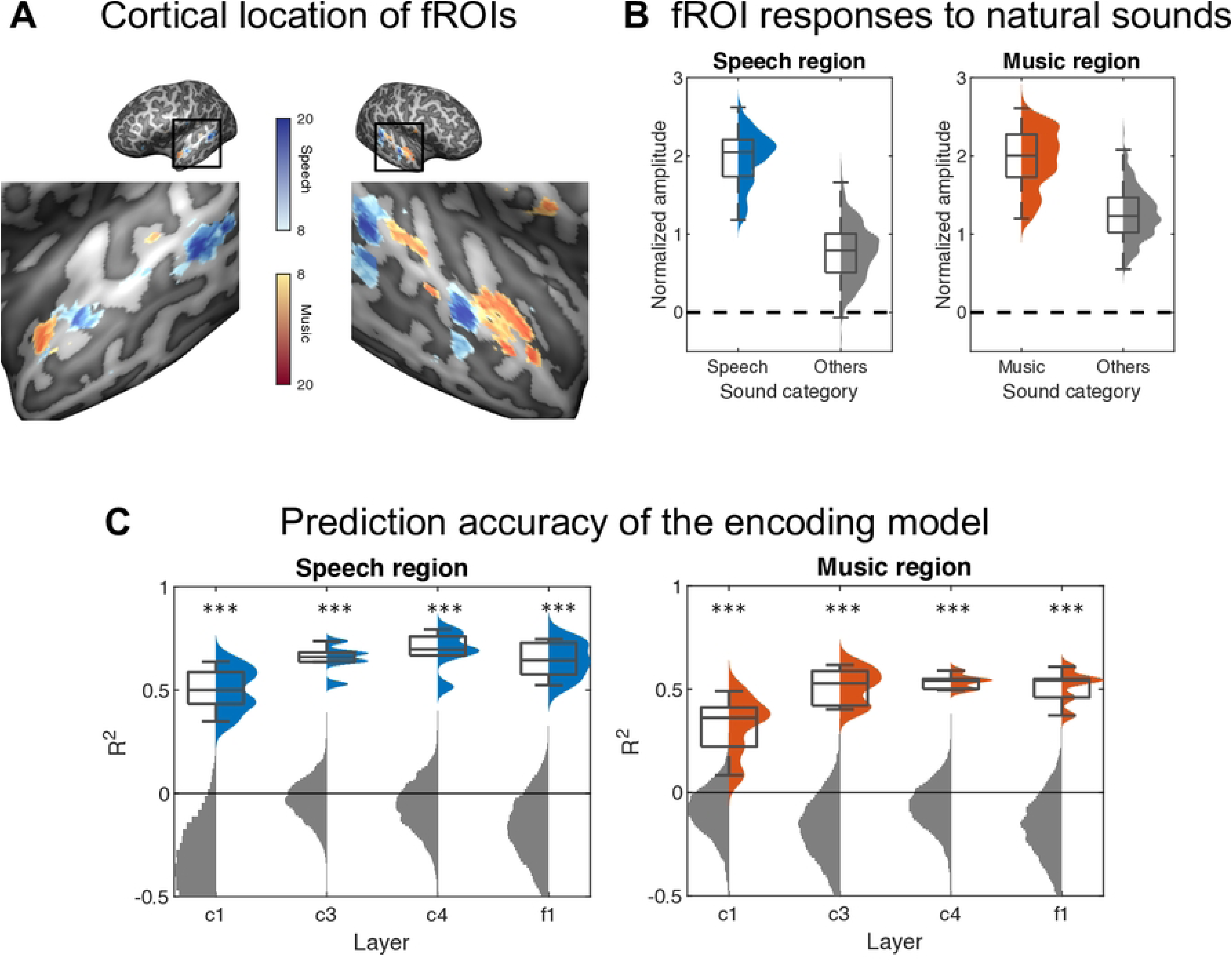
fROI localization and encoding results from a single participant. **(A)** Cortical map of the speech-preferring voxels (blue) and music-preferring voxels (red) that constituted the speech fROI and music fROI, respectively. The color scale represents t-values (denoted sound > all other categories). **(B)** Responses of the speech fROI (left) and music fROI (right) to natural speech (blue) or music (red) and other natural sound categories (gray). Per our fROI definition, the speech fROI showed strong responses to natural speech, and the music fROI showed strong responses to natural music, compared to all other natural sounds. Boxplots show summary statistics across trials. **(C)** Prediction accuracy of the encoding model. The model was used to predict fROI responses to the natural sounds based on ANN layer representations of these sounds. Prediction accuracy was found to be significantly above chance level (permutation-based null distribution, gray) for each layer and fROI. Summary statistics across trials: center, median; lower/upper box limits, first/third quartile; bottom/ top whisker, data within 1.5 interquartile ranges from first and third quartiles, respectively.

### ANN-based sound synthesis

We utilized the fitted encoding model to synthesize sounds with putatively category-relevant features as follows. We first generated 30 white noise samples as starting waveforms. We iteratively increased or decreased their value at each time point to increase the model’s predicted fROI response to the waveform. Each initial waveform underwent this iterative procedure independently for each ANN layer and fROI, resulting in 240 unique synthetic sounds (30 samples × 4 ANN layers × 2 fROIs) for each participant. We refer to sounds synthesized to activate the speech fROI and music ROI respectively as “speech activators” and “music activators”. Fig 3 shows the spectrogram of an exemplary synthetized sound for P2, for each fROI and ANN layer (for other participants, see S2 Fig). Audio files are provided in Supporting information. Auditory inspection of these sounds revealed that they elicit highly unnatural, unintelligible percepts, but with audible differences across layers and sound categories (speech activator vs. music activator). Visual inspection of the spectrograms indicated increasing complexity across layers in spectrotemporal structure, and again noticeable differences across sound categories. For example, speech activators synthesized from layers 3 and 4 exhibited “formant”-like features, whereas music activators synthesized from those layers showed more “tone”-like features.

**Fig 3.**
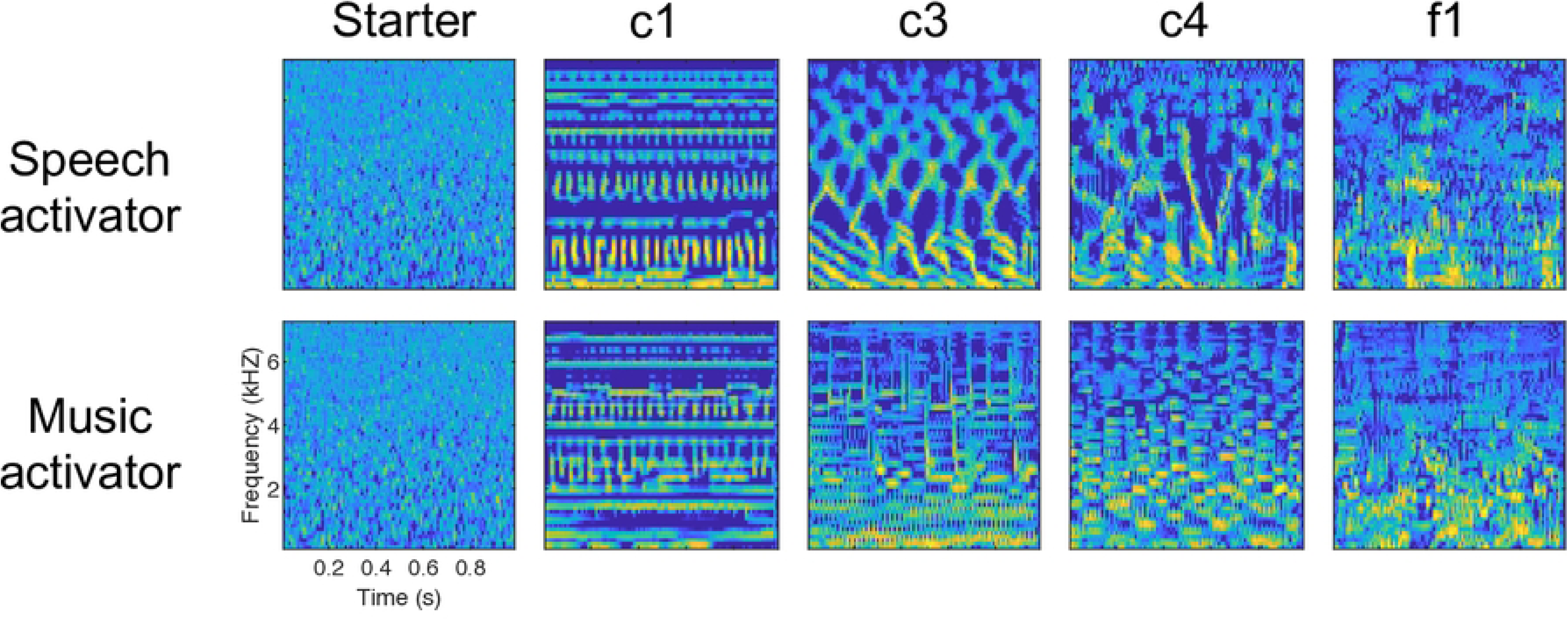
Examples of the ANN-based synthetic sounds. The leftmost column shows the spectrogram of a frozen sample of white noise that served as starting sound waveform for the iterative synthesis procedure. The same sample was used to synthesize one sound that activates the speech fROI (“speech activator”, upper row) and another sound that activates the music fROI (“music activator”, lower row). The other columns represent the ANN layer whose sound-evoked response was used in the synthesis procedure to predict and maximize the fROI response. Data are shown for participant P2; for other participants, see S2 Fig. For audio files, see Supporting audio.

### Representational analysis of synthetized sound features

To characterize the features of the synthesized sounds, we analyzed the representation of these sounds in the ANN using representational similarity analysis [21]. We first tested the categorical nature of the features, by comparing the representations of the synthetic sounds to those of their natural categorical counterparts. We initially focused on layer c4, as sounds synthesized from that layer exhibited the strongest categorical percepts in informal listening tests (for validation, see fMRI and behavioral results). Fig 4A visualizes the representational dissimilarity matrix (RDM) of the c4 representations of all synthetic and natural sounds, after its projection into a two-dimensional space using multidimensional scaling (MDS). The closer two sounds (dots) are in the MDS plot, the more similar are their representations. We observed that synthetic sounds were distant from natural sounds, probably reflecting their unnatural origin. Despite this large representational distance, the synthetic speech activator was closer to the natural speech, and the synthetic music activator was closer to natural music, in line with our informal listening tests. This observation was confirmed by a statistical analysis comparing the distance from speech activator (or music activator) to natural speech (or music) vs. its distance to the other five categories of natural sounds (Fig 4B; speech: p < 1.7 × 10^-91^; music: p < 3.0 × 10^-112^; Bonferroni-corrected across categories). These results indicate that the ANN-based sound synthesis promoted acoustic features that differ from those of natural sounds but distinguish natural sound categories.

**Fig 4.**
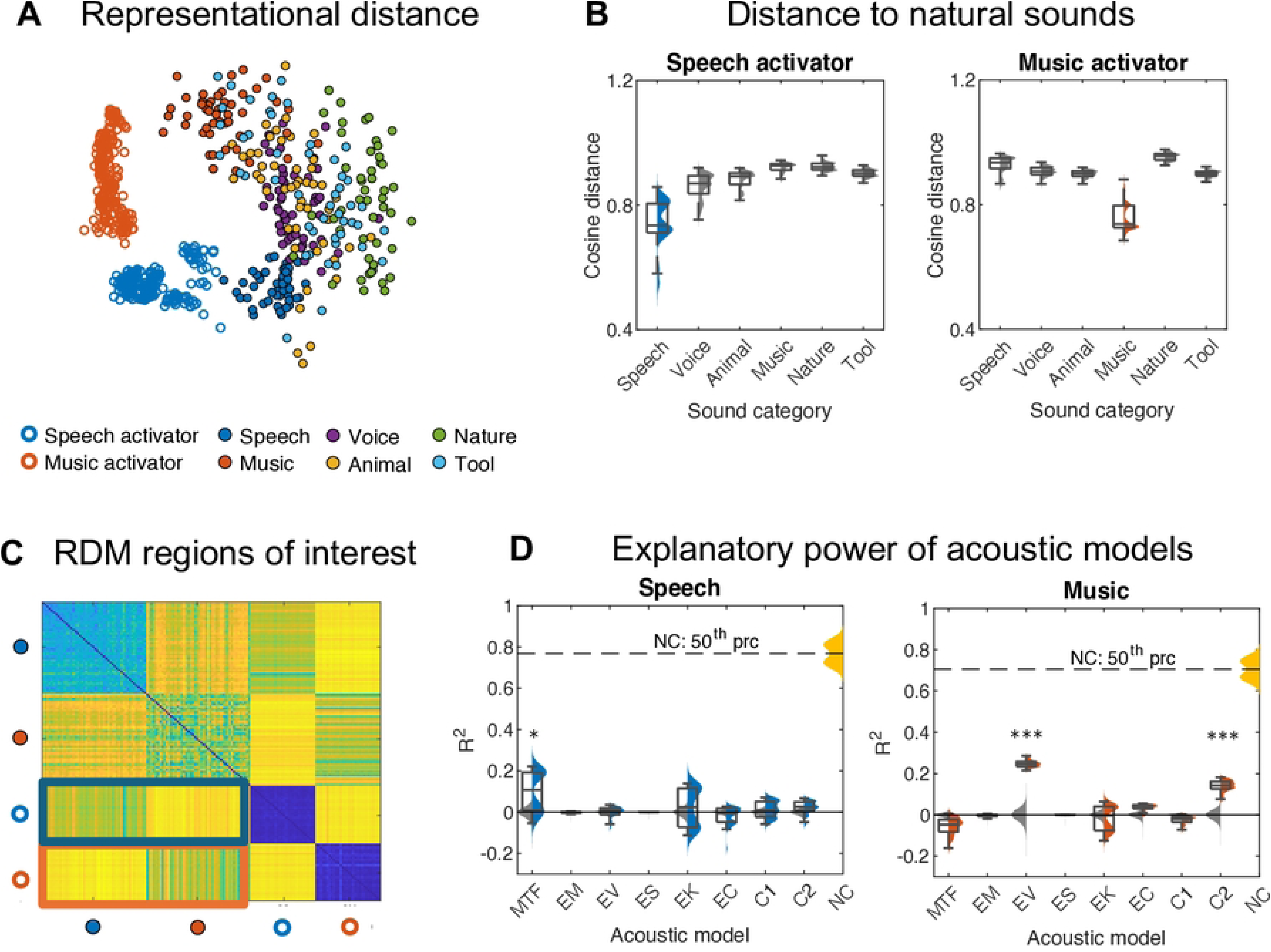
Results from analysis of sound representations in ANN layer 4. **(A)** Two-dimensional (MDS-based) visualization of the representational similarity of the synthetic and natural sounds in c4 (synthetic sounds tailored for participant P2). Each dot represents the representation of a given sound in ANN layer 4, and the distance between dots represents the representational dissimilarity of the sounds. It can be seen that synthetic sounds are distant from natural sounds and the synthetic speech activator and music activator are visually closer to natural speech and music, respectively. **(B)** Representational distance of speech activator (left) and music activator (right) from the six categories of natural sounds in layer 4. In each plot, the colored distribution is significantly lower than the gray distribution. These results confirm our hypothesis that the synthesized sounds possess categorical features. **(C)** RDM regions of interest. The different circles at the margins represent different sounds as labeled in panel A. The blue outline highlights the region of the RDM showing that the speech-activator representation is closer to the natural speech (vs. music) representation in layer 4. The red outline highlights the corresponding region for the music-activator representation. **(D)** Linear regression results quantifying the proportion of the c4-based representational dissimilarity pattern that could be explained by MTF- or texture-based representational dissimilarity patterns, separately for each synthetic sound category / RDM region (speech activator: left plot, blue outline in panel C; music activator: right plot, orange outline in panel C; MTF, modulation transfer function; EM, envelope mean; EV, envelope variance; ES, skewness; EK, kurtosis; EC, envelope correlation; C1 and C2, modulation correlations, see method). Gray distributions are null distributions obtained from 10000 permutations; Orange distributions represent noise ceiling (NC, dashed line, median of NC) across participants. It can be seen that dissimilarity patterns of MTF- and texture-based sound representations could explain a significant, relatively small proportion of the dissimilarity patterns of c4-based speech- and music-activator representations, respectively. Summary statistics across participants: center, median; lower/upper box limits, first/third quartile; bottom/ top whisker, data within 1.5 interquartile ranges from first and third quartiles, respectively. *, P < 0.05; **, P < 0.01; ***, P < 0.001; n.s., non-significant.

We next assessed the uniqueness of the synthetic features by comparing their categorical nature with that of acoustic features known to distinguish sound categories, i.e., those promoted by MTF and acoustic texture models. The MTF represents the spectrotemporal modulation of sounds, while the acoustic texture represents the marginal moments of the amplitude envelope (e.g., mean, EM; variance, EV; skewness, ES; kurtosis, EK) and pairwise correlations across frequency channels (envelope correlations, EC; modulation correlations, C1 and C2, see method). We reasoned that observation of a second-order isomorphism, i.e., a resemblance between the ANN-based RDM and an RDM based on MTF or texture model, would indicate a functional link between the sound features promoted by the models [21]. We focused on the two RDM regions reflecting the categorical nature of the synthetic sounds in the ANN, see Fig 4C: the “speech-activator region” shows the speech-activator representation being closer to the natural speech representation vs. natural music representation, and the “music-activator region” shows the corresponding pattern for the music-activator representation. Using ridge linear regression, we quantified the resemblance between the ANN-based RDM and the MTF- or texture-based RDM, separately for each RDM region and each acoustic model (see Methods). Statistical comparison with chance level (estimated from 10000 permutations) showed that the dissimilarity pattern of MTF-based representations, but not that of texture-based representations, could significantly explain the dissimilarity pattern of the ANN-based representations in the speech-activator region (p = 0.011, corrected across models), and vice versa for the music-activator region (EV, p = 10^-4^, C2, p = 10^-4^, corrected across models). The proportion explained variance was respectively 0.11 and 0.25 in speech-activator region and music-activator region, given a theoretical maximum of 0.78 and 0.70. These results indicate that the categorical features of the synthesized sounds resemble those promoted by MTF and acoustic texture models to only a small degree.

Applying identical analyses to the sounds synthesized from the other ANN layers largely replicated our observations for sounds from layer 4 (S3 and 4 Figs): activator sounds were distant from natural sounds, the speech activator was closer to the natural speech, and the music activator was closer to natural music (compared with other natural sound categories) in all ANN layers. Moreover, only small proportions of the dissimilarity patterns of the ANN-based synthetic sound representations could be explained by the dissimilarity patterns of MTF- or texture-based representations (S1 Table). This indicates that the categorical characteristics reported above were not limited to layer 4, but emerged in all ANN layers tested.

### Effects of synthetized sound features on sound categorization in auditory cortex

To test whether the synthetic sound features influenced sound categorization in the auditory cortex, we presented participants with the synthesized sounds and assessed their effects on speech- and music-fROI responses.

Each participant received their own, individually-tailored set of sounds (60 synthesized from each layer c1, c3, c4, and f1; 24 natural speech, and 24 natural music), except for participant P7 who accidentally was presented with sounds tailored for P6. Fig 5A renders the results of P2, who showed the strongest response differences between speech activator and music activator (see S5 Fig for other participants). We first confirmed that the fROIs showed significant responses to natural sounds as expected (speech fROI: all p < 10^-6^; music fROI: all p < 10^-22^) and the synthetic sounds (speech fROI: all p < 10^-4^; music fROI: all p < 10^-20^) compared to baseline. We further verified that the fROIs responded preferentially to the natural speech vs. music as expected per fROI definition. A 2 × 2 (sound category × fROI) ANOVA confirmed a significant interaction for natural sounds (all participants, p < 10^-6^); the speech fROI responded preferentially to natural speech vs. music (all p < 10^-4^, Bonferroni-corrected), and the music fROI responded preferentially to natural music vs. speech (five participants p < 0.002 Bonferroni-corrected; all p < 0.05 uncorrected; see S5 Fig).

**Fig 5.**
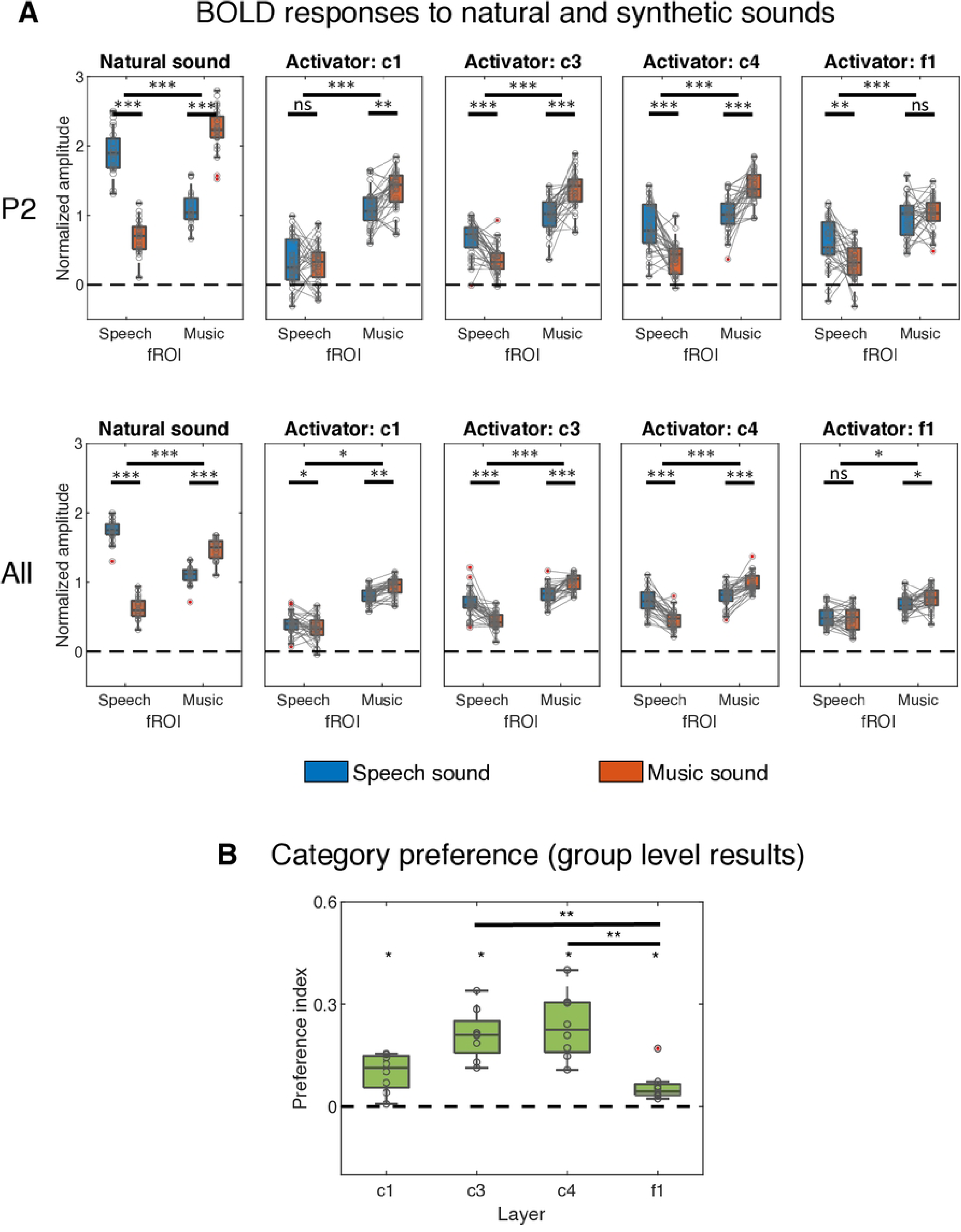
Neural validation of the synthetized sounds. **(A)** The top row shows individual results from participant P2, who showed the strongest response difference between speech activator and music activator (for all other participants, see S5 Fig). In each boxplot, the upper significance marker indicates a significant interaction (sound category × fROI), whereas the lower markers show the simple effect of sound category per fROI, i.e., response difference between speech and music sounds. The solid lines highlight response differences within pairs of sounds synthesized from the same initial noise sample. The interaction between sound categories and the fROIs was significant for natural sounds as expected. Importantly, for synthetic sounds, the same interaction was found, consistently across all layers from which these sounds were synthesized. The bottom row shows results at the group level (N = 8). **(B)** Group results (N = 8). The preference index (PI) quantifies the pairwise response difference between speech activator and music activator (speech fROI: speech activator *minus* music activator; music fROI: music activator *minus* speech activator) averaged across pairs and the two fROIs. PI was significantly above zero for all layers, and significantly higher for sounds synthesized from the middle layers (c3 and c4) vs. sounds synthesized from the later layer (f1). Descriptive statistics across trials (panel A top) or across participants (panel A bottom, panel B): center, median; lower/upper box limits, first/third quartile; bottom/ top whisker, data within 1.5 interquartile ranges from first and third quartiles, respectively. *, P < 0.05; **, P < 0.01; ***, P < 0.001; n.s., non-significant.

After this initial data quality check, we tested our main hypothesis using the same analyses as above, i.e., we assessed whether the fROIs exhibited the category-preference observed above also for the synthetic sounds. Similar to the natural sounds, we found a significant interaction effect (sound category × fROI) for sounds synthetized from the middle layers, c3 and c4. Remarkably, we found this double dissociation reliably in every participant, including the participant who received another’s synthetic sounds (c3: all p < 0.039; c4: all p < 0.044; Bonferroni-corrected across layers; S2 Table 2). We observed this effect also with sounds synthesized from the earlier or later layer, depending on the participant (S5 Fig). Simple-effect analysis confirmed for every participant that the speech fROI preferred the speech activator vs. music activator, and the music fROI preferred the music activator vs. speech activator, from the middle layers (c3 and c4), although this difference was not statistically significant in every participant (S3 Table).

To allow generalizing our individual finding of category preference to the population level, we tested the category preference with group-level statistics. We quantified category preference with a preference index (PI), defined for the speech fROI as the pairwise difference “response to speech activator *minus* response to music activator” (both synthetic sounds generated from the same starting waveform), and vice versa for the music fROI. We pooled PI across all pairs (i.e., starting waveforms) and the two fROIs. As expected, based on our individual double-dissociation results, we found that PI was significantly larger than zero for every layer (Fig 5B; one-tailed Wilcoxon signed-rank tests, Bonferroni-corrected p = 0.016 for all layers). We further observed a main effect of layer on PI (Kruskal-Wallis test, p = 0.001); a post-hoc analysis revealed significantly higher PI in the middle layers than in layer f1 (c3, p = 0.008; c4, Bonferroni-corrected p = 0.003).

In sum, these fMRI results reveal that the synthetic sounds reliably evoked preferential responses in speech and music fROIs qualitatively similar to their natural counterparts. They further show that this category preference is strongest for sounds synthesized from the middle ANN layers. This supports our hypothesis that the synthetic sounds contain features that influence sound categorization in the auditory cortex.

### Effects of synthetized sound features on sound-category perception

To test whether the synthetic sound features influenced sound-category perception, we presented the synthesized sounds to the respective fMRI participants, as well as a group of naïve participants, in a speech/music categorization task with subsequent confidence ratings, and we assessed the effects on behavioral responses.

We presented 240 individually-tailored synthetic sounds (60 synthesized from each layer; for naïve participants, these sounds matched those from a randomly chosen fMRI participant), 30 natural speech, 30 natural music, and 60 control sounds that were synthesized from layer c4 as before but using a “naïve” (i.e., untrained) ANN. On each trial of the task, participants listened to a sound and subsequently categorized it as “speech-like” or “music-like” and rated their confidence in their categorization. “Speech-like” responses to natural speech or speech activator were deemed as “correct” and the same for “music-like” responses to natural music or music activator. Categorization performance was defined as the percentage of all trials on which participants gave a correct response (chance level: 50%; Fig 6a).

**Fig 6.**
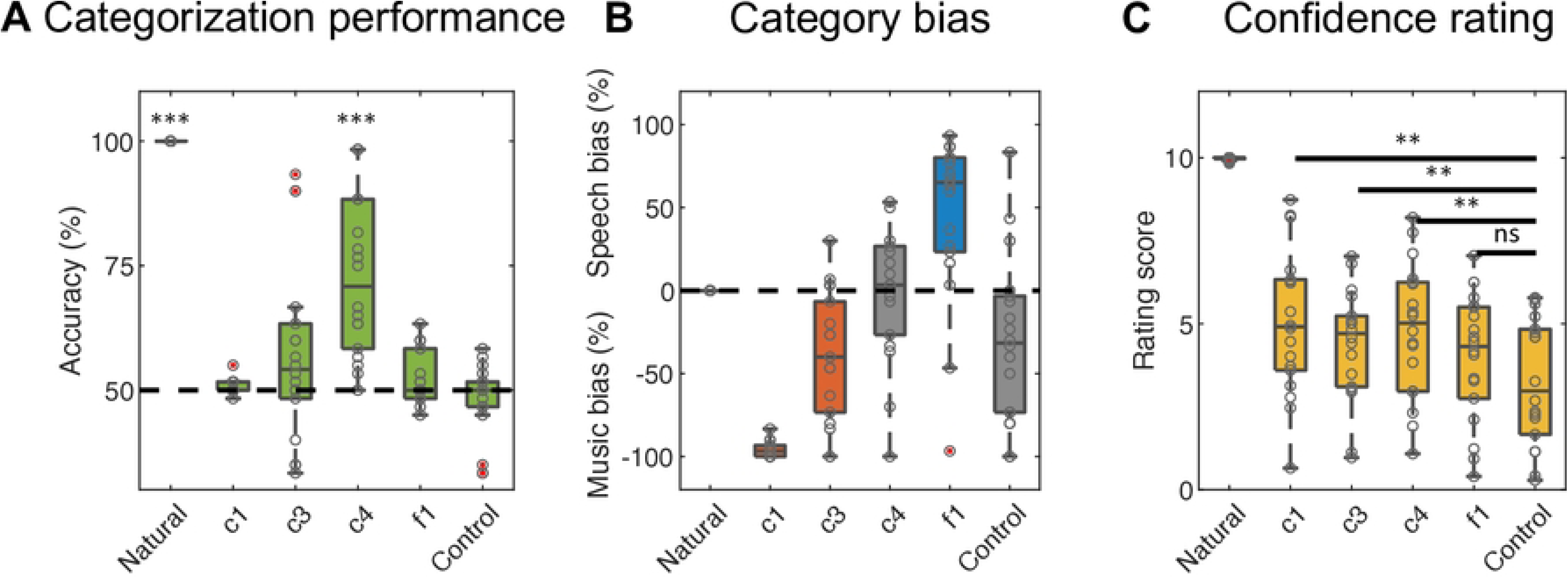
Perceptual validation of the synthetized sounds. **(A)** Categorization performance was defined as the percentage of trials on which participants categorized the sound correctly (“speech-like” response to the natural speech or speech activator, or “music-like” response to the natural music or music activator). Categorization of natural sounds and sounds synthetized from c4 was found to be significantly better than chance level (50%), but not for sounds synthetized from the other layers or a naïve ANN. **(B)** Category bias was computed as the performance difference speech *minus* music; positive values indicate bias to speech, negative values indicate bias to music. The sounds synthesized from c1 and c3 were found to induce significant bias to music (red), whereas sounds synthetized from f1 induced significant bias to speech (blue). Importantly, sounds synthetized from c4 or an untrained ANN induced no such bias (gray). **(C)** Participants’ subjective confidence in their categorization. Participants showed the strongest confidence in their categorization of the natural sounds, and the weakest confidence for the control sounds (synthetized from a naïve ANN). Summary statistics across participants: center, median; lower/upper box limits, first/third quartile; bottom/top whisker, data within 1.5 interquartile ranges from first and third quartiles, respectively. *, P < 0.05; **, P < 0.01; ***, P < 0.001; n.s., non-significant.

We observed no performance difference between fMRI participants vs. naïve participants; therefore, we pooled their data in subsequent analyses. Categorization performance for natural sounds was 100%, consistently across all participants, indicating that they executed the task correctly. In line with our hypothesis, we found that participants categorized synthetic sounds from c4 correctly (median performance: 71%) and significantly above chance (p = 3.6 × 10^-5^). Sounds synthesized from the other layers and control sounds failed to show this effect (p > 0.05, Bonferroni-corrected across ANN layers).

To control for potential category bias, we calculated the performance difference between speech activator vs. music activator. Positive values of this measure indicate a bias to speech; negative values indicate a bias to music (Fig 6b). We found a significant bias toward music for synthetic sounds from c1 (p = 3.6 × 10^-23^; two-tailed t-tests vs. zero, Bonferroni-corrected across ANN layers) and c3 (p = 0.002), and a significant speech bias for synthetic sounds from f1 (p = 0.008). Thus, sounds synthesized from the early layers elicited significantly more music-than speech-like percepts, and vice versa for sounds from the final layer, which might explain our observation of an overall inaccurate categorization of these sounds. We observed no bias for synthetic sounds from c4 (p = 1.000), control sounds (p = 0.096), or natural sounds (p = 1.000), indicating that the accurate categorization of sounds from c4 observed above could not be attributed to bias.

We further explored participants’ subjective confidence in their categorization (Fig 6c). We observed the highest confidence ratings for the natural sounds and the lowest confidence ratings for the control sounds; the latter may be attributed to the fact that these sounds did not incorporate the sound-classification ability of the pre-trained ANN. Confidence ratings for the other synthetic sounds fell in-between, with highest rating for sounds synthesized from layer 4 and layer 1. In statistical support of these observations, we found significantly higher confidence for synthetic sounds from the pre-trained ANN than for the control sounds, for all layers, except the final one (c1, p = 0.001; c3, p = 0.008; c4, p = 0.002; f1, p = 0.077; Bonferroni-corrected across layers).

In sum, these behavioral results validate that the sounds synthetized from layer 4 reliably evoked categorical percepts qualitatively similar with their natural counterparts, in line with our fMRI results above. This further corroborates our hypothesis that these synthetic sounds contain features that influence sound categorization in the human auditory system.

## Discussion

The aim of this study was to identify primary sound features underlying sound categorization in the human auditory system. Using an ANN-based feature-identification approach, we first synthesized novel auditory stimuli to strongly activate cortical regions preferring natural speech or music. We then experimentally confirmed the categorical nature of the features of the synthesized sounds and their relevance for speech- and music categorization in the auditory cortex and perception. We found that the identified categorization-relevant features overlap only partially with those promoted by MTF or texture models. We provide sonified descriptions of the features as audio files (see Supporting information).

Natural auditory environments contain a multitude of sound categories. We focused on two categories that have been consistently shown to undergo distinct processing in the auditory cortex, speech and music. The cortical regions that we observed to preferentially process natural speech or music (speech- and music-fROIs) were widespread across the STG in both hemispheres. Although their exact spatial pattern varied visibly across participants (see S1 Fig), there appeared a tendency for the speech fROI to occupy more posterior portions of STG, whereas music was processed in more anterior regions, including a so-called “pitch region” in the anterolateral extension of Heschl’s gyrus [22]. This aligns with previous studies that have reported the speech fROI and music fROI respectively in regions laterally and anterior to the primary auditory cortex [6,7].

We observed that fMRI responses to natural sounds in the speech- and music ROIs can be predicted from representations of these sounds in the ANN, with average prediction accuracies of up to 0.70 (layer 4−speech fROI). This aligns well with previous fMRI results that have shown prediction of sound-evoked auditory cortical responses with significant, above-chance accuracy [14,15]. Complementary to the previous studies that focused on spatially distributed activation patterns, our results show that the ANN can be utilized to predict also the average activation level within those patterns, which we found to be more suitable for the sound synthesis and yield higher prediction accuracies in pilot data. Overall, our observations confirm that the ANN provides an accurate model for the processing of natural categorical sounds in human auditory cortex [14] and extend previous studies that have proposed analytical models to explain category-preferring auditory cortical responses [6,11]. Importantly, the significant prediction outcomes highlight the utility of the ANN for applications such as the synthesis of sounds to activate specific auditory cortical regions, as discussed below.

Unlike previous fMRI studies that synthesized stimuli by iteratively tuning meaningful, higher-order features (e.g., natural-looking images or words), our study involved the tuning of individual time points of the acoustic waveform. This choice was inspired by previous visual monkey electrophysiology work that tuned individual image pixels, and allowed identifying basic, low-level features within the ANN [18]. Perhaps resulting from this methodological choice, we found the synthesized sounds to have a highly artificial quality, as reflected by our observation of a large difference between these synthesized sounds vs. natural sounds in the ANN. Nevertheless, despite their unnatural appearance, the synthesized sounds exhibited features that distinguish speech from music, as shown by their higher similarity to natural sounds of the same (vs. different) category in the ANN. Based on this, we infer that the synthesized sounds contain categorical features.

We further observed that the categorical features of the synthetized sounds in the ANN are functionally linked with the categorical features promoted by MTF and acoustic texture models, reflecting their shared ability to distinguish sound categories. However, we found the strength of this link to be rather poor compared to noise ceiling. This indicates that the ANN represents sound categories based on features that largely differ from those employed by the other models.

While an analytical specification of the identified features remains difficult, relating them to well-established sound features has revealed two informative insights here: first, the synthetized sounds share some features with corresponding natural sounds of the same category, yet they are composed of mostly other, unnatural features. Second, the categorical features of the synthetic sounds overlap only little with sound features that have been previously associated with categorization (MTF, texture), highlighting their novelty.

Our results identify novel features that are relevant for sound categorization in the human auditory system. Instead of testing a set of pre-defined features hypothesized to be relevant for sound categorization, our study explored previously unknown features that we identified within the feature space of the ANN. As the feature identification was steered by the (predicted) activation level of the category-preferring cortical regions, the resulting features can be inferred to contain information that is primarily processed within those regions or, put differently, constitute the basis of the internal sound representation of those regions. Our fMRI results confirm that the ANN-based synthetic sounds evoke preferential speech and music processing in the auditory cortex that is qualitatively similar to that of natural speech and music, respectively. It should be noted that this observation of preferential responses is non-trivial, given that our sound synthesis targeted the activation level of a given fROI while disregarding the activation levels of other regions. Instead, our fMRI results provide strong confirmatory evidence for the previously reported speech- and music-selectivity of the auditory cortex (for similar reasoning, see Murty et al [15]). Most importantly, these results confirm our hypothesis that the synthetized sounds contain relevant features that determine sound categorization in the auditory cortex.

The results from the behavioral experiment lend further support for this at the perceptual level: participants categorized the sounds synthesized from layer 4 “correctly”, i.e. in line with the preferred category of the cortical region that these sounds were synthesized to activate. Perhaps unsurprisingly, no such effect could be observed with control sounds synthesized in the same way but from a naïve ANN, suggesting that the sound-categorization relevance of the identified features emerged from the sound-classifying ability of the trained ANN.

Another noteworthy side observation is that participants who were stimulated with their own, individually-tailored synthetic sounds, showed no superiority in sound categorization compared to participants who received someone else’s sounds. While this might be attributed to a lack of statistical power, it could also indicate that the identified categorical sound features are rather fundamental and common across participants. More research is needed to address this question.

In sum, our experimental results show that sounds synthetized to activate cortical speech or music regions elicit categorical cortical and perceptual responses that are qualitatively similar to those to natural speech and music. Based on this, we infer that the synthesized sounds can be considered as synthetic speech and synthetic music, and they contain features that play a primary role for sound categorization in the human auditory system.

Regarding layer effects, our results from representational similarity analysis indicate that the synthetic sounds contained categorical information, regardless of the layers that were utilized for their synthesis. Thus, all layers appeared to embed categorical sound features, probably reflecting their ability to jointly classify sounds. However, not every layer’s embedded features are sufficient for sound categorization in the human auditory system: our fMRI and behavioral results consistently show the strongest categorization effect for sounds synthesized from the middle layers, especially layer 4. This suggests that sound features embedded in the ANN layers up to layer 4 contribute to sound categorization in the human auditory system in a cumulative way, whereas subsequent layers may add relatively little to it. Therefore, the sound representation in layer 4 appears to be the most suitable candidate for the internal sound representation in the category-preferring auditory cortex.

In conclusion, speech and music categorization in the human auditory cortex relies on sound features that are captured by an ANN trained for sound classification. These features differ from previously reported MTF- and texture-like acoustic features for sound categorization, and may provide a more accurate description of the internal sound representation of speech- and music-preferring regions in the human auditory cortex. This internal representation might abstract categorical information at a level that is intermediate between those of the low-level acoustic and semantic representations reported previously. Beyond these theoretical insights into sound categorization, our study highlights the utility of combined ANN-FMRI recordings for tailoring sensory stimulation to selectively activate cortical regions. This approach may be useful for non-invasive investigations or clinical treatment of a variety of human brain functions.

## Materials and Methods

### Participants

Eight healthy participants (median age = 29, two males) participated in the fMRI experiment. They also participated in the subsequent behavioral experiment, together with 10 additional healthy participants (median age = 24, four males). All participants reported normal hearing and no history of a hearing disorder or neurological disease and gave informed consent before undergoing the experimental procedures. The experimental procedure was approved by the local ethics committee of the Faculty of Psychology and Neuroscience, Maastricht University (ERCPN #_233_09_02_2021).

### Extraction of fROI responses to natural sounds

#### Auditory stimuli

In the first two fMRI sessions, the auditory stimulation consisted of 288 recordings of natural sounds [23], with 48 sounds from each of the following six semantic categories: speech, (nonspeech) voice, animal cries, musical instruments, nature scenes, and tools. All sounds had a 1-s duration including 10-ms ramps at their onset and offset. They were sampled at 16 kHz, matched for RMS, filtered to correct for non-flatness of the frequency response of the MR-compatible in-ear system (Sensimetrics S14), and presented at a comfortable sound level.

#### fMRI procedure

To establish six unique fMRI runs, the 288 sounds were randomly split into six sound sets with equal, uniform category distributions. Within each set, the sounds were randomly ordered, with the constraint that a given category was never repeated immediately. Sounds were presented in blocks of five, so that each block included five presentations of the same sound. To encourage participants to focus on the sounds, they were instructed to press a button when they detected the target. The latter was defined as a 7-dB level decrease applied to a single, randomly chosen sound repetition in every block. Loudness-change detection performance ranged from 72.5% to 96.1% (mean ± SEM: 84.9 ± 0.03%) across participants given a chance level of 25%, suggesting that all participants paid attention to the auditory stimulation. After every fourth block (or “trial”) a baseline block was presented that was identical to the aforementioned stimulation blocks, except that no sound or task was presented. For participant P1, the stimulation protocol included a larger number of blocks with shorter durations (blocks of four sounds alternated with baseline blocks of half the duration of sound blocks). The aforementioned procedure was conducted twice (i.e., in two fMRI session) with a gap of 1-7 days in-between. The fMRI data collected in these two sessions served the definition of individual fROIs and encoding models (explained in detail below).

MRI data were acquired with a 3T scanner (Siemens) at Scannexus (https://scannexus.nl/). Anatomical T1-weighted images were acquired using a modified magnetization-prepared rapid-acquisition gradient-echo sequence (number of slices = 192, TR = 2250 ms, TE = 2.21 ms, flip angle = 9°, voxel size = 1 mm^3^ isotropic, GRAPPA acceleration factor = 3). BOLD signal changes were recorded using a modified gradient-echo echo-planar imaging (EPI) sequence isotropic (number of slices = 33 without gap, TR = 2400 ms, TA = 1000 ms, TE = 30 ms, flip angle = 82°, voxel size = 2 mm^3^ isotropic, GRAPPA acceleration factor = 3). Functional image acquisition and sound presentation alternated to ensure clear audibility of the sounds in the absence of scanner noise. Functional slices were centered on the Sylvian fissures to cover the temporal lobes. All functional and anatomical images were acquired in anterior-posterior phase encoding direction. To enable subsequent correction for EPI distortions, an additional pair of EPI volumes with reversed phase-encoding direction was acquired before the first functional run [24].

#### fMRI data preprocessing

Imaging data were preprocessed with BrianVoyager 22.4 (Brain Innovation). Functional data were processed in four steps including slice scan-time correction (using cubic spline interpolation), 3D motion correction (using trilinear/sinc interpolation), temporal high-pass filtering (cutoff frequency: 7 cycles per run), and EPI distortion correction (using the TOPUP method). The anatomical and functional data from all fMRI sessions were spatially coregistered and normalized to Talairach space.

The fMRI signal of each voxel was z-scored along the time course within each run and delayed by one TR relative to the stimulation protocol to compensate for the hemodynamic delay. Functional volumes acquired in response to the four sound repetitions within a given block were averaged to obtain a single, time-averaged response measure for that block (the volume acquired immediately after the first sound was rejected to reduce carryover effects from the preceding block). The same approach was applied to baseline blocks to obtain a time-averaged baseline response.

#### fROI selection

The fROIs were selected in three steps [6]: first, voxels showing significant sound-evoked responses were identified for each session by comparing sound blocks vs. baseline blocks (p < 0.001, one-tailed t-test). Second, from these sound-responsive voxels, the ones showing a response profile that was most reliable across sessions were selected. Reliability *r* was assessed as follows:

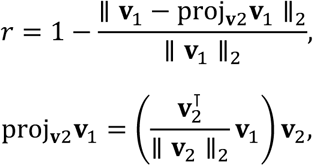

where vector **v**_1_ is the response of a given voxel to the 288 sounds in the first session, and **v**_2_ is the same voxel’s response to the same sounds in the second session. Only voxels fulfilling a reliability criterion of r > 0.3 were retained (except for P1, we retained top 10% voxels). Finally, to identify a speech fROI, the retained voxels were submitted to statistical analysis contrasting speech blocks vs. no-speech sound blocks (one-tailed t-test) and the 5% of voxels showing the largest t-value were selected (all p < 0.001). Correspondingly, a music fROI was identified using the contrast music blocks vs. no-music sound blocks. Voxels within a given fROI were pooled to obtain a single, spatially averaged response measure.

### Extraction of ANN responses to sounds

We focused on the VGGish [20], an ANN that has been pre-trained to classify 10-s YouTube audio tracks with a set of 527 labels from the Audioset ontology [25]. The VGGish has a relatively simple architecture and has been proven successful in predicting human auditory cortex response patterns to natural sounds [14]. The VGGish takes audio segments of 0.975-s duration (converted to a stabilized log-mel spectrogram) as input and feeds it through an architecture consisting of an input layer, four convolutional blocks (convolution, ReLU, MaxPooling), and three fully connected blocks (full connected, ReLU) that progressively reduce dimensionality, resulting in a 128-dimensional semantic embedding (output layer). We fed the 288 natural sounds into the VGGish and extracted representations of these sounds from its convolutional layers and fully connected layers as follows: after generating a cochleagram of each sound, we passed it through the VGGish, and extracted the response of each model unit, from each layer. The convolutional layers represented the sound along three dimensions: time, frequency, and kernel. To facilitate prediction of the time-averaged fROI responses, we averaged the time dimension yielding a time-averaged vector whose length (4096 points) equaled the product of the number of frequency points and number of kernels. A vector of equal length was directly extracted from the fully connected layers (i.e., without time dimension). Thus, this procedure resulted in eight sets of fixed-size (4096 points) sound representations (conv1, conv2, conv3_1, conv3_2, conv4_1, conv4_2, fc1_1, fc1_2). Considering the computational load and the time limit of the fRMI recording, we limited the validation experiments on four layers covering the early to late processing stages of the ANN architecture: conv1, conv3_1, conv4_1 and fc1_1.

### Encoding models: mapping ANN responses onto fROI responses

We next trained an encoding model to map the ANN representations of the sounds onto the measured fROI responses to these sounds, separately for each fROI and layer. We modeled each fROI’s time-averaged response to a given sound as a linear combination of the 4096 points defining that sound’s time-averaged representation in a given layer. The model was fitted using regularized linear regression and a six-fold cross-validation procedure. The six unique sound sets from the six fMRI runs were split into five training sets (used to fit the model) and one test set (used to evaluate the accuracy of the prediction of the fitted model). Each set contained responses to 48 unique sounds and was z-scored independently. On each iteration of the procedure, a different sound set was defined as test set. Given that the number of regressors (4096) exceeded the number of training sounds (240), L2-regularized regression was used. Introducing the L2-penalty on the weights results in a closed-form solution to the regression problem, which is similar with the least-squares regression normal equation:

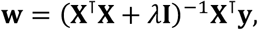

where w was a 4096-point column vector containing the regression weights, y was a 240-point column vector representing the response variable (i.e., the fROI‘s response to each sound), X was a 240 × 4096 matrix representing the predictor variable (i.e., the layer-specific representation of each sound), and I was the 4096 × 4096 identity matrix. The ridge parameter was optimized within the range *λ* = 10^2^, 10^1^, …, 10^-6^, 10^-7^. The value yielding the most accurate predictions (maximum mean predictive variance) was retained. The predictive variance R^2^ was estimated as 1 – SSEtest/SSTtest, where SSEtest represents the sum of the squared error of the trained model’s prediction of the test-set data and SSTtest represents the total sum of squares of the test-set data. To obtain an empirical chance level, we computed 1000 permutations across the test-set sounds for each split. The same permutations were applied to the corresponding sounds in the training set. The aforementioned procedure was conducted separately for each fROI × layer combination (2 × 4).

### Sound synthesis

We utilized the VGGish and the fitted encoding model to synthesize sounds via a gradient-ascent optimization approach. The procedure started with 30 white noise samples, the RMS level of which was matched to the natural sounds. The noise was fed into the VGGish, its representation was extracted from each layer, and the fROI response to this noise was predicted using the fitted encoding model. The value of each time point of the noise waveform was adjusted in the direction that increased the predicted fROI response, using a cost function based on the negation of the predicted fROI response and the total variation between directly neighboring time points (the latter served to reduce acoustic transients). The learning rate was set to a value of 0.001. The gradient of the cost function was calculated across 700 iterations using the function built in TensorFlow 2.13 (www.tensorflow.org), after which the optimized sounds showing the minimum costs were chosen. In each iteration, the novel sound from the previous iteration was temporally shifted in random direction (forward or backward) with a maximum of 1 ms to generate the adversarial component; the latter served to promote the synthesis of sounds that elicit predicted fROI response levels unaffected by such small shifts. The above procedure was conducted separately for each fROI × layer combination (2 × 4), each time starting with the same set of 30 (frozen) white noise samples. Overall, this resulted in a set of 240 (2 × 4 × 30) unique synthetic sounds.

### Representational analysis of synthetized sounds

Representational dissimilarity analysis (RSA) [21] was applied to representations of the sounds in the ANN, the MTF model, and the sound texture model. VGGish representations of the synthetic sounds were extracted as described above (see section *Extraction of ANN responses to sounds*). The MTF was computed using NSLtools [26] as follows: first, the sound cochleagram was computed, which consisted of the time-varying ouput (temporal resolution = 8 ms) of 128 cochlear filters with log2 spaced frequencies (179–7246 Hz). Then, the magnitude of the spectrotemporal modulations (scale from 0.25 to 8 cycles/octave; unsigned rate from 2 to 32 Hz) in each channel of the cochleagram representation was assessed and averaged across time points. The sound texture was analyzed using the Sound Texture Synthesis Toolbox v.1.7 [27]. The texture model considered here includes four separate components measuring different summary statistics of the time-varying amplitude envelopes of the cochleagram (32 frequency bands evenly spaced on an ERB-rate scale from 20 Hz to 10 kHz), or of the modulation analysis specific to each of the frequency bands (six modulation filters with center frequencies log-spaced between 30 and 100 Hz): (1) the marginal statistics of the band-specific envelopes (mean, EM; variance, EV; skewness, ES; kurtosis, EK); (2) the pairwise correlations C between band-specific amplitude envelopes (EC); (3) the pairwise correlation between the modulation analysis of each frequency band at a constant modulation filter frequency (C1 correlations); and (4) the correlation between the modulation analysis of the same frequency band at adjacent modulation filter frequencies (C2 correlations). For each type of representation, between-stimulus cosine distances were computed using a customized function in MATLAB, and all parameters were given equal weight.

We used ridge linear regression to predict the group-averaged RDM of the ANN from the group-averaged RDMs of the acoustic models (cross-validated across both stimuli and participants). For the stimuli, we focused on natural speech, natural music, speech activator, and music activator. More specifically, we focused on the RDM regions that revealed the categorical nature of the synthetic sounds, i.e., where the speech activator was close to natural speech and distant to natural music; and the reverse for the music activator, see Fig 4C. All the sounds were split into a training set and a test set each including half of the sounds of each category, and the sample was split into a training group and test group consisting respectively of six and two participants. All possible combinations of sound splits and sample splits were tested, resulting in 56 folds. The predictive variance *R^2^* was estimated as described above (see section on encoding models). Each fold included the estimation of a null distribution (10,000 row × column permutations of the RDM) as well as a cross-validated measure of noise ceiling that captured the maximum predictable variance in the group-averaged ANN distances. The latter was estimated as 1 – SSDtest-train/SSTtest, where SSDtest-train represents the sum of the squared differences between the group-averaged ANN distances in the test vs. training set.

Statistical significance was assessed with a one-sided Mantel’s test comparing the observed *R^2^* values with the permutation-based null distributions followed by a maximum-statistics correction for multiple comparisons [28].

### Neural validation of synthetized sounds

The synthetized sounds were validated at the neural level in a third fMRI session that matched the two preceding fMRI sessions in all regards (participant, stimulation protocol, fMRI protocol, task, and fMRI data preprocessing), except for the following: the stimulation included the 240 synthesized sounds (2 categories × 4 layers × 30 samples) and only 48 natural sounds (24 speech, 24 music). The synthetic sounds were preprocessed identically to the natural sounds (see above). The 24 natural speech and music sounds were the ones observed to elicit the strongest speech-fROI and music-fROI response in the two preceding fMRI sessions. The analysis focused on fMRI responses extracted from the fROIs defined in the preceding fMRI sessions.

### Behavioral validation of synthetized sounds

The behavioral experiment used the same study design as the fMRI experiment, additionally including a synthetic-sound condition that served to control for potential bias in participants’ category ratings of synthetic sounds. For that condition, synthetic sounds were generated as before, but with the difference that a random-weights VGGish with Kaiming He weights initialization was used and only layer c4 was considered. Thus, the design contained twelve conditions: 2 fROIs × 4 layers, plus 2 natural sounds, and 2 control sounds from the random VGGish. Participants who had undergone the fMRI part received the same sounds as in the fMRI experiment (see section above), except that 30 (instead of 24) unique natural sounds were presented to match the number of corresponding unique synthetic sounds. For each of the other, naïve participants, the sounds were the same as for a pseudorandomly chosen fMRI participant. The stimuli were presented via a sound card and headphones (Sennheiser, HD650) at a sound level of 65 dB SPL.

On each trial of the speech/music categorization and rating task, participants were presented with a sound and instructed to categorize the stimulus as either ‘speech-like’ or ‘music-like’ and rate their confidence in this categorization on an 11-point Likert scale (0 = not confident at all; 10 = totally confident). Responses were entered via a GUI that was continuously displayed throughout the experiment. The position (left or right) of the ‘speech-like’ and ‘music-like’ response buttons on the interface was counterbalanced across participants. After participants entered their rating, the next trial started. Trials were presented in blocks of 72, each including six unique sounds from each condition and lasting approximately seven minutes. In total, five blocks were presented, resulting in an overall number of 360 unique trials (30 per condition).

## Supporting information

**S1 Fig.** fROI localization and encoding results for all participants, except for participant P2 who is shown in the corresponding main Fig. 2. For details, see main Fig. 2.

**S2 Fig.** Examples of the ANN-based synthetic sounds for all participants, except for participant P2 who is shown in the corresponding main Fig 3. For details, see main Fig 3.

**S3 Fig.** Representational distance of the speech activator (left column) and the music activator (right column) from the six categories of natural sounds for ANN layers c1, c3, and f1 (rows). For layer c4 and details, see the corresponding main Fig. 4B. The synthetic sounds were significantly more similar to their natural counterparts than to natural sounds from other categories (colored distribution < gray distribution; p < 10^-9^) regardless of the layer from which the sounds were synthesized, except for music synthetized from f1.

**S4 Fig.** Linear regression results quantifying the proportion of the ANN layer-based representational dissimilarity pattern that could be explained by MTF- or texture-based representational dissimilarity patterns, separately for each synthetic sound category / RDM region (columns). Results are shown for ANN layers c1, c3, and f1 (rows). For layer c4 results and further details, see the corresponding main Fig 4D. For inferential statistics, see S1 Table.

**S5 Fig.** Neural validation of the synthetized sounds for all participants, except for participant P2 who is shown in the corresponding main Fig 4A. For details, see main Fig 4. For inferential statistics of each participant, see S2 and 3 Tables.

**S1 Table.** Statistical results from ridge linear regression analysis of RDM regions. Proportion explained variance (median *R^2^*value across cross-validation folds) and associated p-value (in parentheses, permutation-based) are shown for each acoustic model (rows) and synthetic sound / RDM region as represented in layer c1, c3, c4, and f1 (shown respectively in columns 1 and 2, columns 3 and 4, columns 5 and 6, and columns 7 and 8). P-values were adjusted using a maximum-statistics correction for multiple comparisons across acoustic models.

**S2 Table.** Statistical p-values associated with the sound category × fROI interaction, for each participant and sound type (natural sounds and synthetic sounds from each ANN layer). P-values for synthetic sounds were Bonferroni-corrected across layers.

**S3 Table.** Statistical p-values associated with simple effects of sound category (in speech fROI: speech activator > music activator; in music fROI: music activator > speech activator), for each participant and sound type (natural sounds and synthetic sounds from each ANN layer). P-values were Bonferroni-corrected across layers, the uncorrected p-values are shown in parentheses.

**S1 Audio Files.** Examples of synthesized sound.

## Acknowledgement

We thank J. Hewett, M. Leonard, and B. Sorger for assistance with data collection.

## Author contributions

L.X. and L.R. conceptualized the study. L.X., E.F. and L.R. developed the methodology. L.X. wrote the software. L.X., E.F. and L.R. validated the results. L.X. conducted the formal analysis and investigation. E.F. and L.R. provided resources. L.X. curated the data. L.X. and L.R. wrote the original draft of the manuscript. L.X., E.F. and L.R. wrote, reviewed and edited the manuscript. E.F. and L.R. supervised and administered the project and acquired funding.

